# An adipocyte light-Opsin 3 pathway regulates the circadian clock and energy balance

**DOI:** 10.1101/368373

**Authors:** Shruti Vemaraju, Gowri Nayak, Ethan D. Buhr, Yoshinobu Odaka, Kevin X. Zhang, Julie A Mocko, April N. Smith, Brian A. Upton, Jesse J. Zhan, Vishnupriya J. Borra, Elise Bernhard, Kazutoshi Murakami, Minh-Thanh Nguyen, Shannon A. Gordon, Gang Wu, Robert Schmidt, Xue Mei, Nathan T. Petts, Matthew Batie, Sujata Rao, John B. Hogenesch, Takahisa Nakamura, Russell N. Van Gelder, Richard A. Lang

## Abstract

Almost all life forms can detect and decode light information for adaptive advantage. Examples include the visual system, where photoreceptor signals are processed into virtual images, and the circadian system, where light entrains a physiological clock. Here we describe a pathway in mice that employs encephalopsin (OPN3, a 480 nm light responsive opsin) to mediate light responses in murine adipocytes. The adipocyte light-OPN3 pathway regulates neonatal growth in mice and is required for at least three important functions including (1) photoentrainment of a local circadian clock, (2) extracellular matrix deposition, and (3) regulation of mitochondrial content and the proportion of “brite” adipocytes. Furthermore, we show that the light-OPN3 pathway is required for normal levels of uncoupling protein 1 (UCP1) in white and brown adipose tissue. Consequently, neonatal *Opn3* germ-line and adipocyte-conditional null mice show a reduced ability to maintain their body temperature under cold stress. This was also observed in wild-type mice deprived of blue light. We hypothesize that the adipocyte light-OPN3 pathway provides a dynamically responsive, circadian clock-integrated mechanism for regulating adipocyte function and in turn directing metabolism to thermogenesis rather than anabolism. These data indicate an important role for peripheral light sensing in mammals and may have broad implications for human health given the unnatural lighting conditions in which we live.

---

During breeding of mice homozygous for the *Opn3^lacz^* null allele (Fig. S1a), we noted a growth differential in neonates. Assessment of body weight from the day of birth (P1) to P15 revealed that *Opn3^lacz/lacz^* homozygotes grew larger than control littermates (Fig. 1a, b). Homozygotes were, on average, 1.3 gm heavier than controls at P15 (Fig. 1a). Crown-to-rump measurements indicated that *Opn3* null mice were also longer than normal (Fig. 1a, inset chart). At P23, the size difference was obvious (Fig. 1b). Over many litters, the average increase in weight for neonatal *Opn3^lacz/lacz^* mice was 9.1% (Fig. 1c).

**Figure 1.**
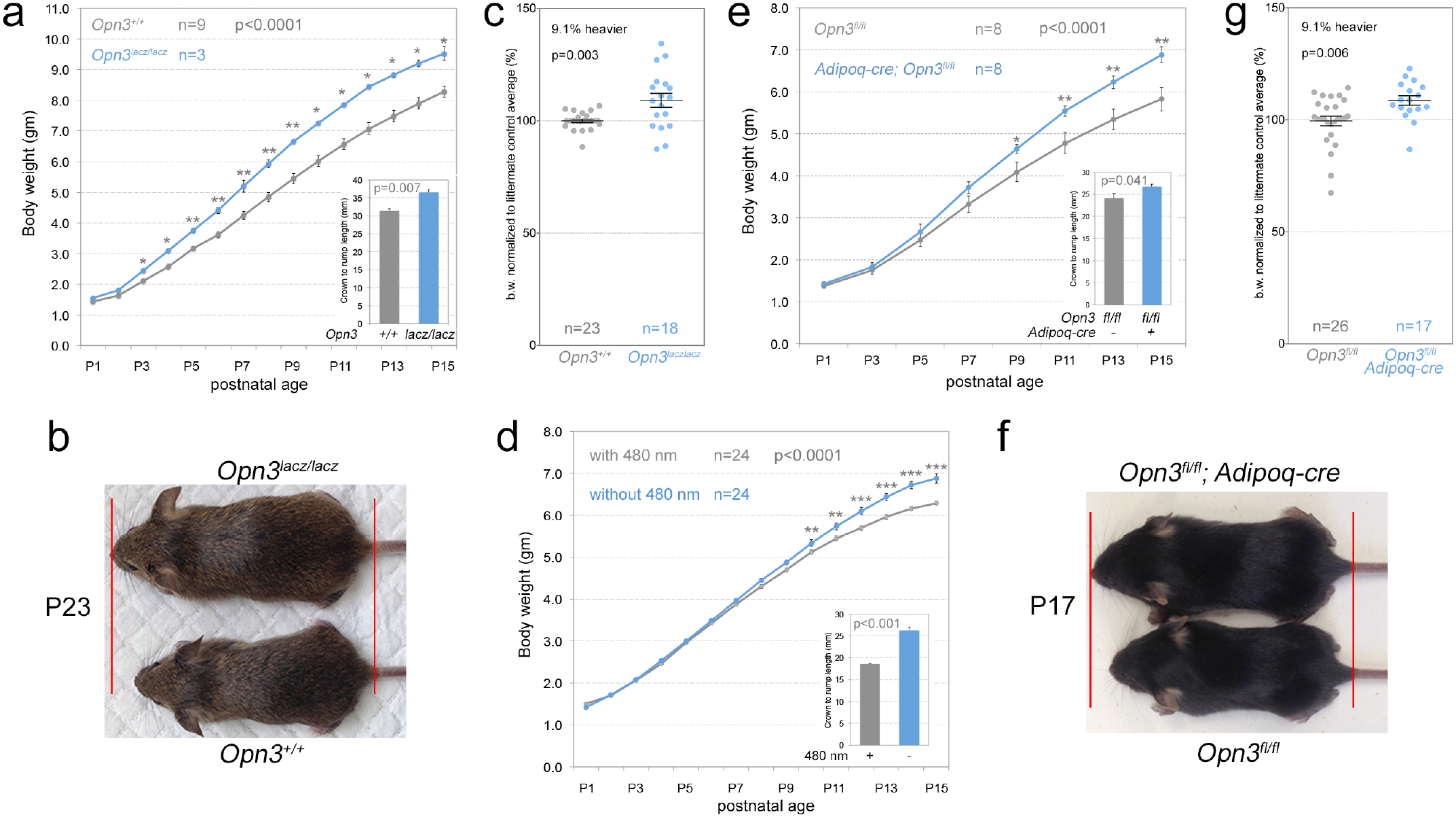
*Opn3* and light-dependent regulation of neonatal growth. (a) P1-P15 growth chart for *Opn3^lacz/lacz^* null and wild-type mice showing that the null mice are larger according to weight. Chart inset shows crown-to-rump length at P15. (b) *Opn3^lacz/lacz^* versus WT mouse size at P23. (c) Scatter plot showing *Opn3^lacz/lacz^* null mouse weight at P16 as a proportion of wild-type controls. They are on average, 9.1% heavier. (d) P1-P15 growth chart for C57BL/6 mice raised from birth with and without 480 nm centred light (λ_max_ for OPN3) showing that the absence of 480 nm light results in larger mice according to weight. Chart inset shows crown-to-rump length at P15. (e) P1-15 growth chart for control *Opn3^fl/fl^* and *Adipoq-cre; Opn3^fl/fl^* mice. Conditional null mice are larger than normal. Chart inset shows crown-to-rump length at P15. (f) *Opn3^fl/fl^* versus *Adipoq-cre;; Opn3^fl/fl^* mouse size at P17. (g) Scatter plot showing *Adipoq-cre;; Opn3^fl/fl^* conditional null mouse weight at P16 as a proportion of wild-type controls. They are on average, 9.1% heavier. All data in this figure are presented as mean ± s.e.m. n is sample size. Statistical significance for growth time-courses (a, e, d) was assessed by two-way ANOVA and p-value indicated on top of chart. All single time-point comparisons were performed with *t*-test *, *P*<0.05, **, *P*<0.01, \*\*\**P*<0.001. In these data, no sex differences were apparent.

OPN3 has the conserved amino acids that qualify it as a ciliary opsin^1^,^2^. Vertebrate OPN3 also forms a functional photopigment when expressed *in vitro*^3,4^. However, it remained possible that the large size of mutant mice had little to do with the light sensing function of OPN3. To assess this we raised C57BL/6J mice (that are generally smaller than the mixed background mice shown in Fig. 1a) in normal 12L:12D lighting (12 hr light, 12 hr dark) that either included or excluded the blue, 480 nm centered wavelengths that stimulate OPN3^3^. This showed that “minus blue” resulted in elevated mouse weight (Fig. 1d) and length (Fig. 1d, inset chart), providing a phenocopy of OPN3 loss. Since melanopsin (OPN4) also absorbs in the 480 nm band, we assessed *Opn4* null mice but did not detect a neonatal overgrowth (Fig. S1). Combined, these data raised the interesting possibility that an OPN3-dependent light response normally influences neonatal growth in a mammal.

*Opn3* is expressed in adipose tissue in mice^5^ and humans^6^ (Fig. S2g, h) raising the possibility that OPN3 activity in adipocytes could explain the neonatal growth phenotype. We performed conditional deletion of a *loxP*-flanked *Opn3* allele (*Opn3^fl^*, Fig. S1) with the adipocyte-specific *Adipoq-cre*^7^. Assessment of body weight and crown-to-rump length (Fig. 1e) showed that *Adipoq-cre;; Opn3^fl/fl^* mice were larger than normal (Fig. 1f), mimicking the phenotype of the *Opn3* germ-line null and selective loss of blue light. Over many litters, the average increase in weight of neonatal *Adipoq-cre; Opn3^fl/fl^* mice was 9.1% (Fig. 1g) identical to that observed for the germ-line null (Fig. 1c). Thus, conditional deletion identified adipocytes as a key site of OPN3 function.

There are many different adipose tissue depots in mice and three different types of adipocytes (white, brown and “brite” or “beige”^8^). No reliable antibodies for murine OPN3 are presently available. To determine the expression pattern of *Opn3* in adipocytes we took advantage of the *Opn3^lacz^* allele (Fig. S1). The interscapular adipose tissue (iAT) depot comprises interscapular subcutaneous white adipose tissue (iscWAT) and interscapular brown adipose tissue (iBAT). Xgal labelling of cryosections from P16 iAT showed no background labelling in control (Fig. 2a) but intense labelling in the *Opn3^lacz/lacz^* iscWAT (Fig. 2b). In iBAT, Xgal labelled cells were not readily apparent at low magnification (Fig. 2b) but at higher magnification and bright transillumination, a subset of expressing brown adipocytes was detected (Fig. 2d). Labelled cells were not present in control iBAT (Fig. 2c). P16 inguinal WAT (inWAT) from control mice was Xgal negative (Fig. 2e). Unlike in the adult, P16 inWAT has a high content of “brite” adipocytes (Fig. 2e, f, BrAd). These do not appear to be Xgal labelled in *Opn3^lacz/lacz^* mice. By contrast, large, unilocular adipocytes from *Opn3^lacz/lacz^* mice were Xgal positive (Fig. 2e-h). For confirmation of this *Opn3* expression pattern, we used the *Opn3^cre^* allele(Fig. S1) to convert the tdTomato reporter *Ai14* (Fig. 2i-k). Cryosections showed that in iAT, almost all adipocytes within the iscWAT were positive (Fig. 2i, k). In iBAT, a subset of brown adipocytes was positive (Fig. 2j, k). Finally, we also used an *Opn3-eGFP* reporter that is based on a bacterial artificial chromosome transgene (GENSAT #030727-UCD). In an assessment of inWAT, iscWAT and iBAT, this reporter confirmed expression of *Opn3* in the majority of white adipocytes and a subset of brown adipocytes in iBAT (Fig. S2). Finally, the GTEx database showed that in human subcutaneous and omental adipose tissue, OPN3 was expressed at low-to-middling levels of 2.9 and 3.1 tags per million, respectively (Fig. S2). These data indicate that human adipose tissues express *OPN3*.

**Figure 2.**
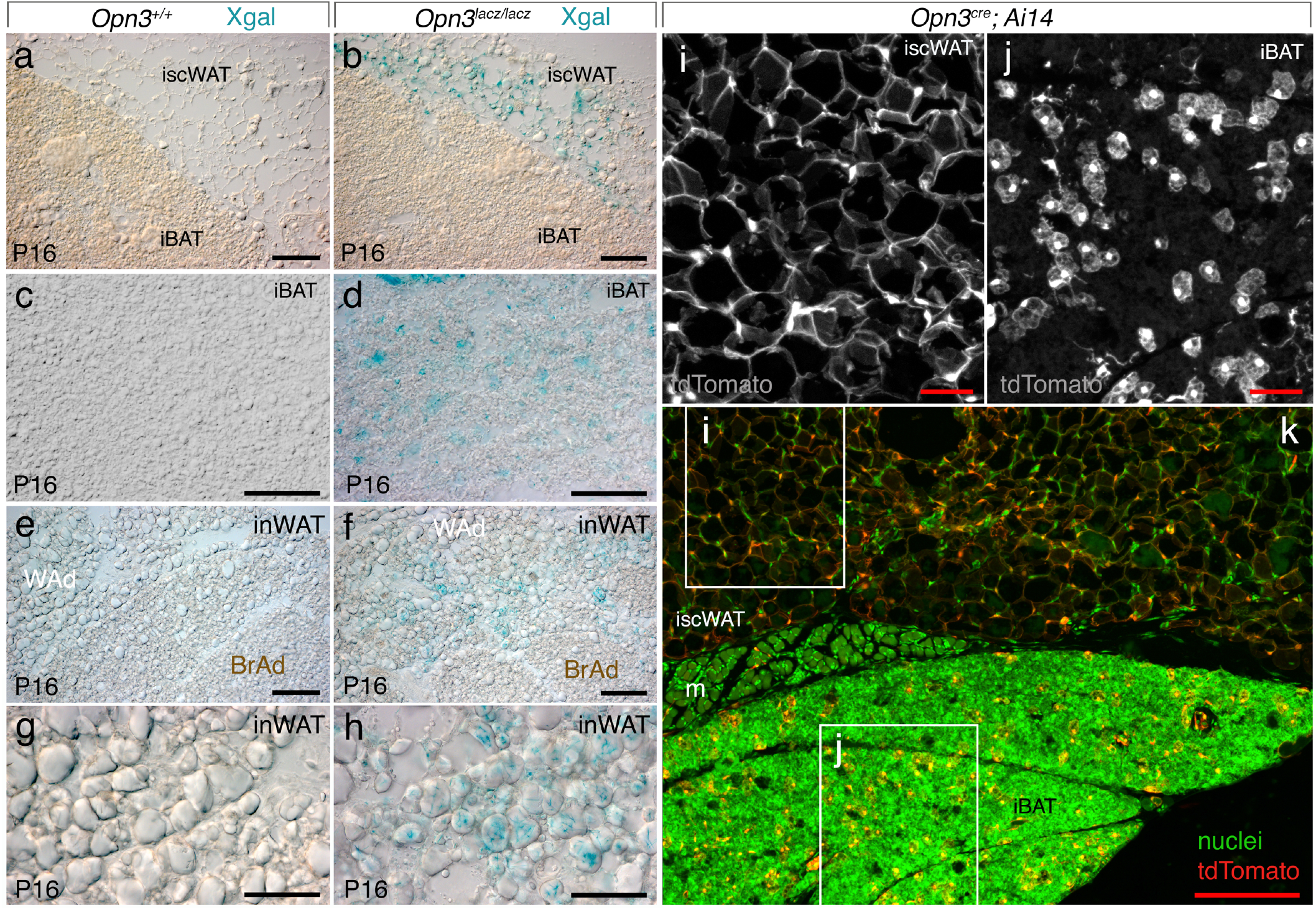
Expression of *Opn3* in iAT and inWAT. (a-d) Xgal labelled wild type (a, c) and *Opn3^lacz/lacz^* (b, d) cryosections of interscapular adipose tissue (iAT) including interscapular subcutaneous white adipose tissue (a, b, iscWAT) and interscapular brown adipose tissue (a-d, iBAT) at P16. (e-h) Xgal labelled wild type (e, g) and *Opn3^lacz/lacz^* (f, h) cryosections of inguinal white adipose tissue including white (WAd) and “brite” (BrAd) adipocytes. (i-k) Detection of tdTomato (red, greyscale) in *Opn3^cre^;; Ai14* mice for iAT showing positive cells in iscWAT (i) and iBAT (j). iscWAT and iBAT are separated by a leaflet of muscle (m) that is visible in some sections. Labelling of nuclei with Hoechst33258 is presented in green. In (a,b,e,f,k) scale bars are 100 µm. In (c,d,g,h,i,j) scale bars are 50 µm.

A major function for ‘non-visual’ opsins is the light entrainment of circadian clocks;; both melanopsin (OPN4) and neuropsin (OPN5) function in this capacity^9,10,11,12^. To determine if OPN3 influenced the circadian clock, we first analyzed adipose tissue for circadian rhythmicity by monitoring bioluminescence from tissue explanted from PER2::LUC mice^13,14^. iscWAT and inWAT from wild-type mice showed clear rhythmicity *ex vivo* (Fig 3a-c). In contrast, the same tissues from *Opn3-*null mice showed markedly reduced circadian amplitude (Fig 3a, b, red traces, c, similar to that previously described in *Opn3-*null retina^12^). Furthermore, *in vivo* qPCR assessment of the transcripts for clock genes *Per1-3, Bmal1, Npas2, Nr1d1/2* and *Rora* in control *Opn3^fl/fl^* and experimental *Opn3^fl/fl^; Adipoq-cre* mice at two different circadian time-points (ZT1 and ZT17, Fig. 3d) showed significant down-regulation of *Per1-3* at ZT1 and of *Per2, Per3, Npas2, and Nr1d1/2* at ZT17. This is consistent with the reduced amplitude of PER2::LUC rhythm in acute *ex vivo* assessment (Fig. 3a-c) and suggests that OPN3 function affects the adipose circadian clock *in vivo*.

**Figure 3.**
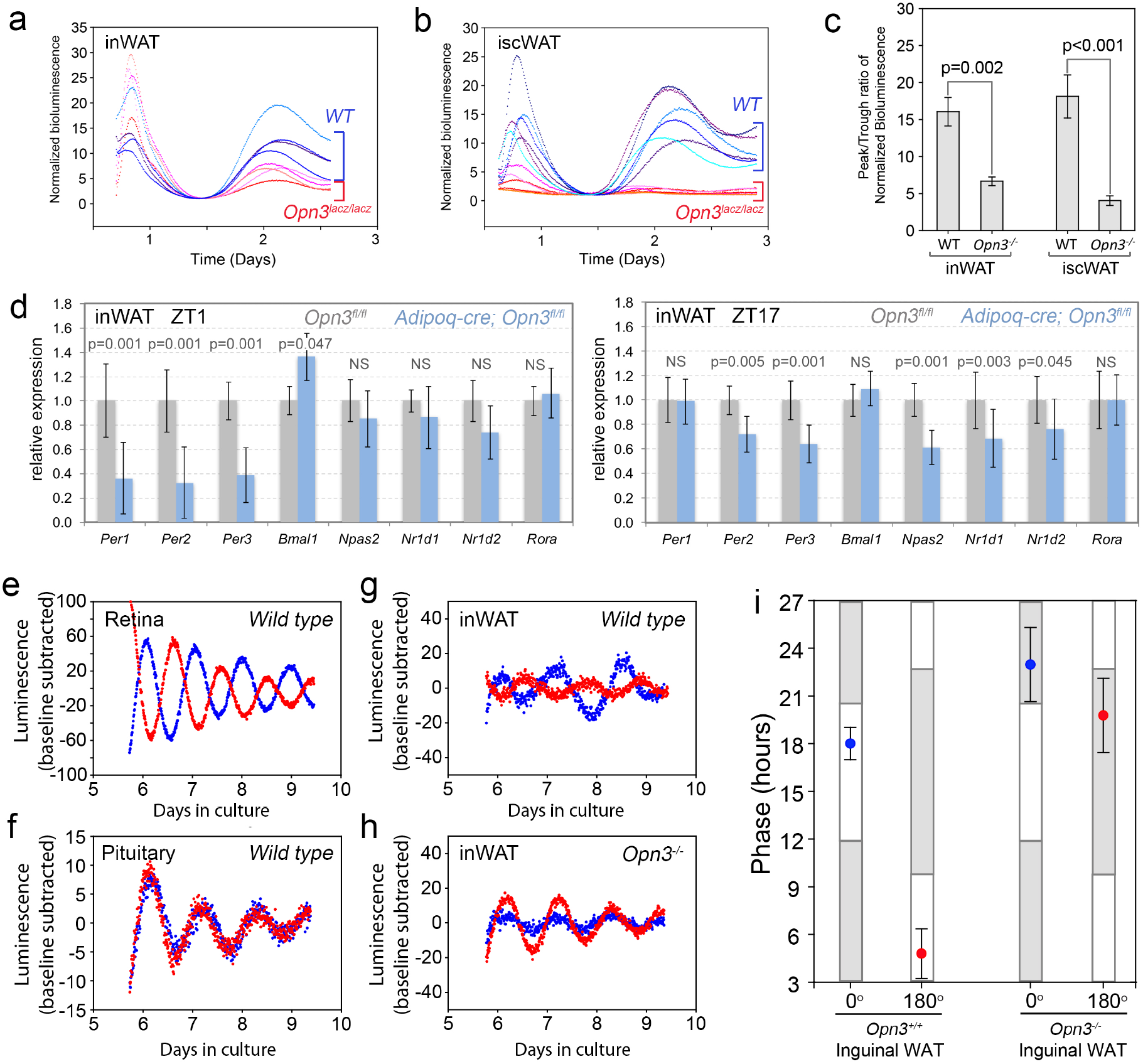
*Opn3* is required for light-dependent circadian clock function in WAT. (a, b) PER2::LUC circadian clock reporter oscillations in inWAT (a) and iscWAT (b) immediately after explant. In both tissues, clock oscillation in the *Opn3* null shows low amplitude as indicated by the peak-trough ratio (c). (a-c) n=6 for each genotype and tissue. Comparison between genotypes in c were performed with Mann-Whitney Rank Sum test and p-value indicated on chart. (d) qPCR analysis for circadian clock genes *Per1-3, Bmal1, Npas2, Nr1d1/2, and Rora* in inguinal WAT of *Opn3^fl/fl^* and *Opn3^fl/fl^; Adipoq-cre* mice at P16 for ZT1 (left) and ZT17 (right). The expression level of many clock genes is reduced. n=3 for each genotype and time-point. Statistical significance calculated using fixed pairwise randomization reallocation test. (e) Representative traces from photoentrainment assay for retina showing PER2::LUC luminescence readout for samples exposed to LD versus DL. (f) as in (e) but for pituitary. Retina photoentrains, pituitary does not. (g) Opposing phases of PER2::LUC readout from inWAT indicates photoentrainment. (h) Loss of opposing PER2::LUC phases in *Opn3* null inWAT photoentrainment. (i) Aggregation of phase data for control (n=6) and *Opn3* null (n=5) inWAT photoentrainment assays indicating a requirement for *Opn3*. Data are presented as mean ± s.e.m.

To determine if OPN3 might be functioning in a phototransduction pathway in this tissue, we assessed whether the circadian clock of adipose tissue could be photoentrained *in vitro* in a manner similar to retina and cornea^12^. Photoentrainment assays monitor the real-time oscillation of the clock gene reporter PER2::LUC^13^ after a period of four days in which the tissue is exposed to an LD cycle^11^. Opposing phases for two explants that experience LD versus DL indicates photoentrainment and light responsiveness^11^. Wild-type retinal tissue showed clear photoentrainment in this assay (Fig. 3e). Tissues that experience LD versus DL and do not photoentrain (like pituitary, Fig. 3f) show coincident PER2::LUC phases. While iscWAT PER2::LUC oscillations were not detectable after six days in culture, inWAT from *Per2::Luc* mice showed low amplitude oscillations that demonstrated clearly opposing phases and thus light-dependent clock entrainment (Fig. 3g). Conversely, inWAT from *Opn3* null *Per2::Luc* mice showed coincident phases (Fig. 3h) indicating that OPN3 is required for light-dependent clock entrainment in adipose tissue. Plotting the average PER2::LUC peak from multiple inWAT photoentrainment assays (Fig. 3i) confirms that OPN3 is required for photoentrainment of inWAT. These results indicate that OPN3 is intimately involved in both free-running maintenance and light-responsiveness of the circadian oscillator in adipose tissue.

To further address the function of OPN3, we performed a microarray-based transcriptome analysis on P16 control and *Opn3* germ-line null mice. Tissues for this analysis were harvested on P16 at ZT1. The control and *Opn3* null mice used in this experiment showed no significant weight difference. This choice was designed to minimize the identification of pathways secondary to *Opn3* loss-of-function. We harvested two tissues (iAT and inWAT) that express *Opn3* and one, liver, that does not. Using the AltAnalyze suite^15,16^, we identified differentially regulated transcripts that fell into functional clusters and pathways. Clustering based on significant Z-scores for WikiPathway models is noted (Fig. S3a-c) but this information is also developed into a more detailed overall schematic (Fig. S3d-h). In addition, subsets of this schematic and additional Opn3-dependent transcriptome changes are shown in Figs. 4-6. In these schematics, each box represents a transcript where red and blue colour coding indicates up- or down-regulation, respectively. With a few noted exceptions, all *Opn3*-dependent transcript regulation is significant to p<0.05. The *Opn3* transcript showed the highest negative fold change in *Opn3* null iAT (3.0 fold down, p=1.5×10^−4^) and inWAT (5.6 fold down, p=7.6×10^−3^) but was not significantly changed in liver where expression levels are very low. Changes in some transcripts of interest were validated by qPCR (Fig. S4). Overall, transcriptome analysis indicated that OPN3 activity was required for normal regulation of metabolism, as evidence by deregulation of lipid, glucose and energy generation pathways. Consistent with a role for OPN3 in regulating adipose circadian clocks, we detected one clock gene, *Npas2*, that was significantly changed in iAT (Fig. S3, S4a).

Transcriptome analysis for both iAT and inWAT revealed remarkable clusters of upregulated extracellular matrix (ECM) transcripts, their integrin receptors and ECM remodelling enzymes (Fig. 4a, b, S4a). For iAT, most of the clustered ECM components are collagens (Fig. 4a), whereas for inWAT (Fig. 4b), there is variety including fibrillins, a fibronectin, a laminin as well as *Col14a1* and *Thrombospondin 3*, the only two regulated transcripts in common with the iAT ECM cluster. Immunodetection of Collagen 6 in iAT and inWAT from *Opn3* adipocyte conditional null mice at P16 (Fig. 4c and d, square) revealed a robust up-regulation in both iBAT (Fig. 4e-h) and inWAT (Fig. 4i-l). This validates transcriptome data but because this is an adipocyte conditional deletion, also shows that *Opn3* has a crucial adipocyte-specific function. In *Opn3* germ-line null mice at P16, an up-regulation of ECM was not obvious by immunofluorescence, perhaps because for each individual transcript, up-regulation was modest (1.1-1.3 fold). However, when we assessed *Opn3* germ-line null mice at 7 months of age, we observed a robust up-regulation of COL6 in both iBAT (Fig. 4m-p) and inWAT (Fig. 4q-t). This suggests that in germ-line *Opn3* null mice, there may be a more subtle, progressive accumulation of COL6, perhaps due to non-adipocyte functions of OPN3. It is known that the ECM of adipocytes, and in particular Collagen 6^17^, is crucial for normal metabolic function^18,19,20^. Adipose tissue fibrosis, a marked accumulation of ECM components, is also associated with metabolic disruption^17^.

The ECM clusters in iAT and inWAT include many transcripts that normally show circadian rhythmicity in expression level according to the CircaDB database^21^. Remarkably, 7 of 9 ECM transcripts in iAT show rhythmic expression (Fig. 4a). These data are also consistent with findings from circadian clock mutant mice. For example, *Clock* mutant mice show a very similar ECM transcript clustering in cardiomyocytes^22^. This reinforces the suggestion that one function of OPN3 is to regulate the adipose circadian clock.

**Figure 4.**
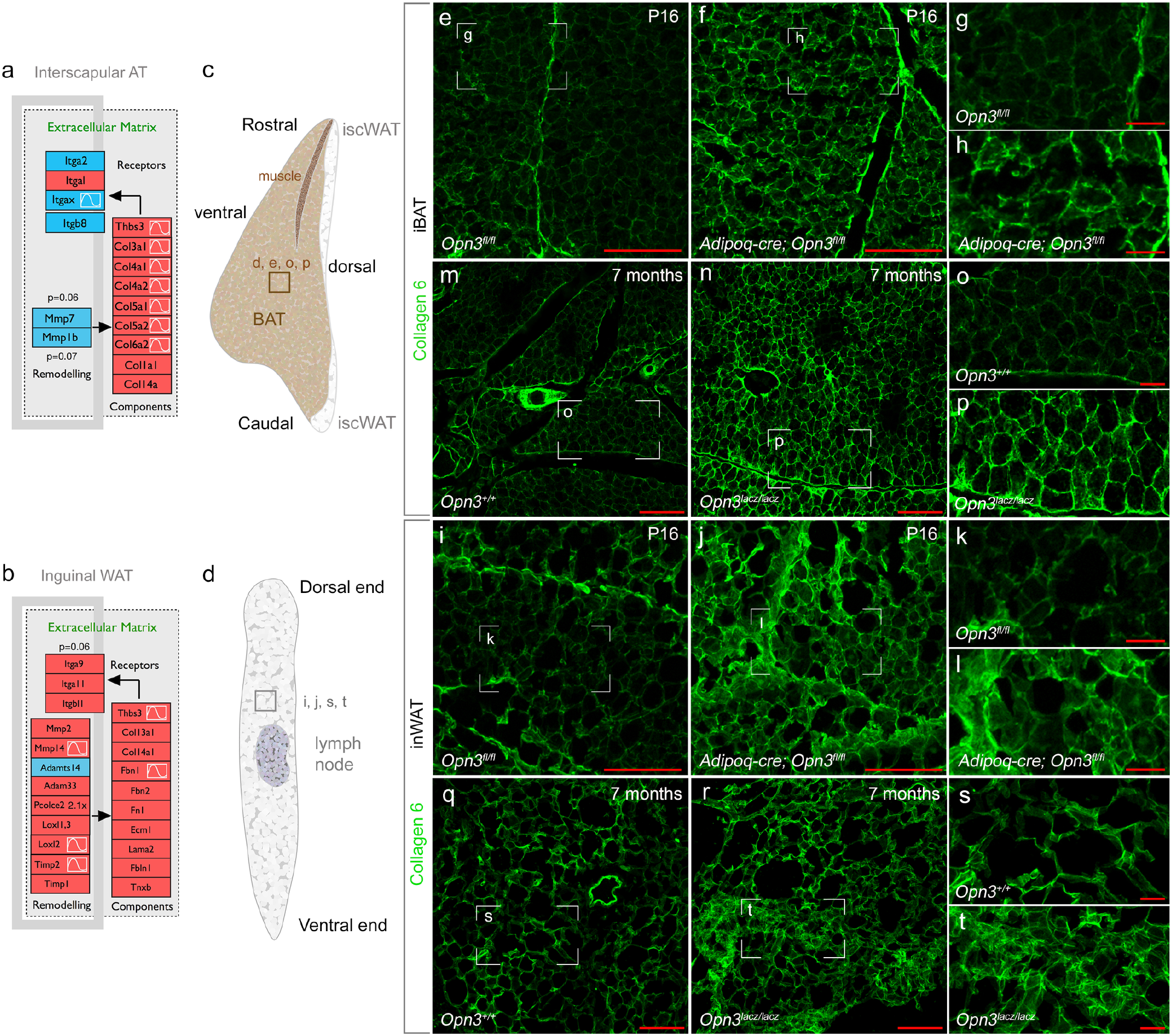
*Opn3* null mouse iAT and inWAT extracellular matrix deregulation. (a, b) Schematic of clustered *Opn3*-dependent transcript changes in ECM signalling pathways for iAT (a) and inWAT (b). Red and blue colour coding indicates up- and down-regulated transcripts, respectively. The sine-wave symbol indicates that, according to data within CircaDB, expression of this transcript is diurnally rhythmic in this tissue type. Schematics are modified versions of Wikipathway models for the Focal Adhesion-PI3K-mTOR Signaling pathway (WP2841). (c, d) Diagrams indicting the regions of iAT and inWAT cryosections in which images for (e-t) were generated. (e-t) Labelling for Collagen 6 (COL6) at P16 in control and *Opn3^fl/fl^; Adipoq-cre* mice (e-h, iBAT and i-l, inWAT) and at 7 months in control and *Opn3^lacz/lacz^* germ-line null mice (m-p, iBAT and q-t, inWAT). (g,h,k,l,o,p,s,t) are magnified versions of the regions indicated. Scale bars in red, 100 µm (e,f,m,n,i,j,q,r), 25 µm (g,h,o,p,k,l,s,t).

In inWAT, *Opn3*-dependent, differentially regulated transcripts cluster within the PPAR pathway (Fig. 5a). This pathway regulates adipocyte size as well as lipid metabolism and energy generation^23,24^. Many PPAR pathway genes are regulated by the circadian clock^23^ and consistent with this and *Opn3*-dependent adipose entrainment (Fig. 3), 9 of 12 regulated PPAR pathway transcripts show rhythmic expression in adipose tissue according to the CircadB database^21^. An assessment of adipocyte size in inWAT of *Opn3* germ-line and adipocyte conditional null mice at P16 (Fig. 5c, d) revealed that both showed significantly elevated adipocyte size. Neonatal WAT is a mixture of white and “brite” adipocytes and normally has a significantly higher proportion of “brite” adipocytes than adult WAT depots^25,26^. Hematoxylin staining of inWAT sections showed that both the *Opn3* germ line (Fig. 5e,f) and adipocyte conditional nulls (Fig. 5g,h) have a lower proportion of the smaller, “brite” adipocytes, as would be suggested by adipocyte size assessment.

**Figure 5.**
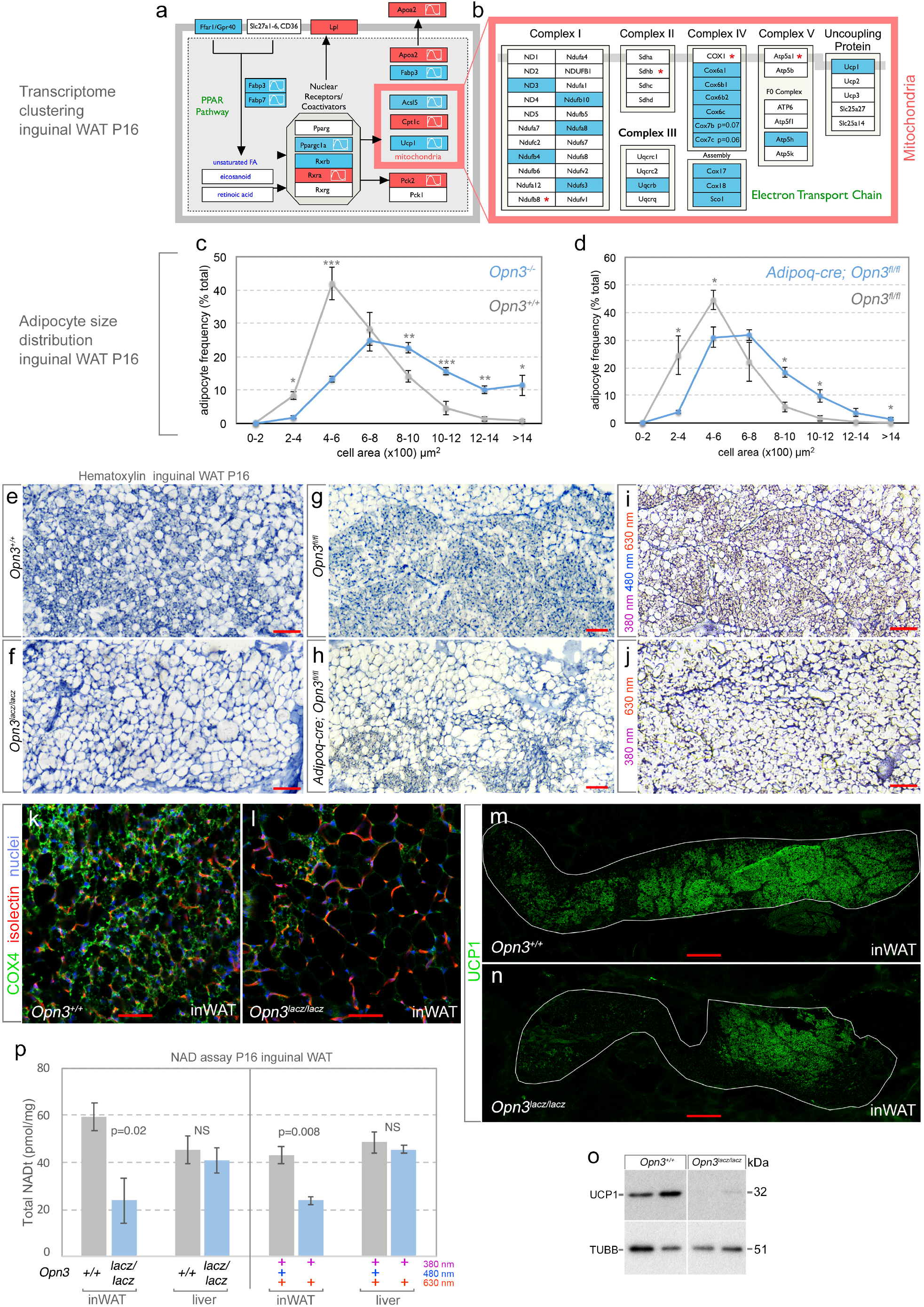
*Opn3* null and “minus blue” reared mouse inWAT phenotype. (a, b) Schematic of clustered *Opn3*-dependent transcript changes in the PPAR pathway (a) and in the electron transport chain of mitochondria (b). Red and blue colour coding indicates up- and down-regulated transcripts, respectively. The sine-wave symbol indicates that, according to data within CircaDB, expression of this transcript is diurnally rhythmic in this tissue type. Schematics are modified versions of Wikipathway models for PPAR Signaling Pathway (WP2316, (a)), and for the electron Transport Chain (WP295, (b)). (c, d) Adipocyte size distribution in inWAT comparing control and *Opn3^la^cz/lacz* (c) as well as control *Opn3^fl/fl^* and *Adipoq-cre; Opn3^fl/fl^*(d) at P16. Data are presented as mean ± s.e.m. n=3 for each genotype. Direct comparisons between genotypes at each interval were performed with Student’s T-test \**P*<0.05, \*\**P*<0.01, \*\*\**P*<0.001. (e-j) Hematoxylin staining of histological sections of P16 inWAT from *Opn3^+/+^* (e), *Opn3^lacz/lacz^* (f), *Opn3^fl/fl^* (g), *Adiopq-cre; Opn3^fl/fl^* (h), full spectrum (380 nm, 480 nm, 630 nm) reared (i), and “minus blue” (380 nm, 630 nm) reared (j) mice. In (f, h, j) diminished numbers of “brite” adipocytes and larger average cell size is apparent. (k, l) Labelling for the electron transport chain marker COX4 (green) in P16 control and *Opn3* null mice. Both reduced COX4 labelling and larger adipocyte size is apparent in the *Opn3^lacz/lacz^* tissue. Counter-labelling is isolectin for vasculature (red) and Hoechst33258 for nuclei (blue). (m, n) UCP1 (green) labelling of inWAT in control (m) and *Opn3^lacz/lacz^*(n) germ-line null animals at P16. Boundary of inWAT is outlined in white. (o) Immunoblot detecting UCP1 and β-tubulin (TUBB) in P16 inguinal WAT from *Opn3^+/+^* and *Opn3^lacz/lacz^* mice. UCP1 levels are low in the *Opn3* null tissue. (p) Total NAD levels in inguinal WAT and liver for P16 *Opn3^+/+^* and *Opn3^lacz/lacz^* mice (left chart, n=4) or for mice reared either in “full spectrum” or “minus blue” lighting (right chart, n=3). p values calculated using Student’s T-test. In both the *Opn3* null and “minus blue” conditions, NAD level are low in inWAT. Scale bars in red (e, f, i, j, k, l) 50 µm, (g, h) 100 µm (m, n) 500 µm.

Consistent with this, inWAT from the *Opn3* null showed a striking cluster of 17 electron transport chain (ETC) transcripts, all of which were down-regulated (Fig. 5b). Confirming a lower mitochondrial content, the ETC components COX4 (Fig 5k, l) and UCP1 (Fig. 5m,n) were both detected at a lower level by immunofluorescence in P16 *Opn3* null inWAT compared with the control. In whole inWAT sections, UCP1 immunofluorescence was widespread in the control, and more restricted in the *Opn3* null (Fig. 5m,n). Low levels of UCP1 in P16 *Opn3* null inWAT were confirmed by immunoblotting (Fig. 5o).

We have shown that, like *Opn3* loss-of-function, “minus blue” lighting causes neonatal mice to overgrow (Fig. 1). To determine whether this phenocopy extended to tissue changes observed in the *Opn3* null, we performed hematoxylin staining of histological sections and found the lower “brite” adipocyte content (Fig. 5i,j) characteristic of both the *Opn3* germ line and conditional nulls. Finally, we assessed inWAT from *Opn3* null and “minus blue” reared mice at P16 for total NAD content as this essential coenzyme mediator of electron transport provides a measure of mitochondrial content and function^27^. This quantification showed that in neither case was there a change in liver NAD (Fig. 5p). However, in both the *Opn3* null and “minus blue” reared mice, NAD levels in inWAT were low (Fig. 5p). Combined, these data indicate that a neonatal light-OPN3 pathway influences the “beigeness” and mitochondrial function of inWAT.

Prompted by the effects of *Opn3* deficiency on mitochondria in inWAT and a modest cluster of ETC genes in iAT (Fig. 6a), we assessed mitochondrial status in P16 iBAT using immunoblot for ETC components and transmission electron microscopy (TEM). Immunoblotting for the ETC components ATP5A (Complex V), COX1 (Complex IV), SDHB (Complex II), NDUFB8 (Complex I) and UCP1 revealed some variability in the presence of SDHB in the *Opn3* null but a consistently low level of both NDUFB8 and UCP1 (Fig. 6b). An immunoblot of P16 mice raised in “minus blue” lighting revealed that similarly, NDUFB8 and UCP1 were at lower than normal levels (Fig. 6c). At P28, mitochondrial morphology in *Opn3* null iBAT was often abnormal with a disorganized pattern of cristae (Fig. 6d). Transcriptome analysis clustering also revealed that four TRP (Transient receptor potential cation channels) family genes were regulated and that Leptin, an adipokine, was 1.9 fold reduced (Fig. 6a). The latter was validated by the low Leptin levels detected in serum of *Opn3* null mice (Fig. S4).

**Figure 6.**
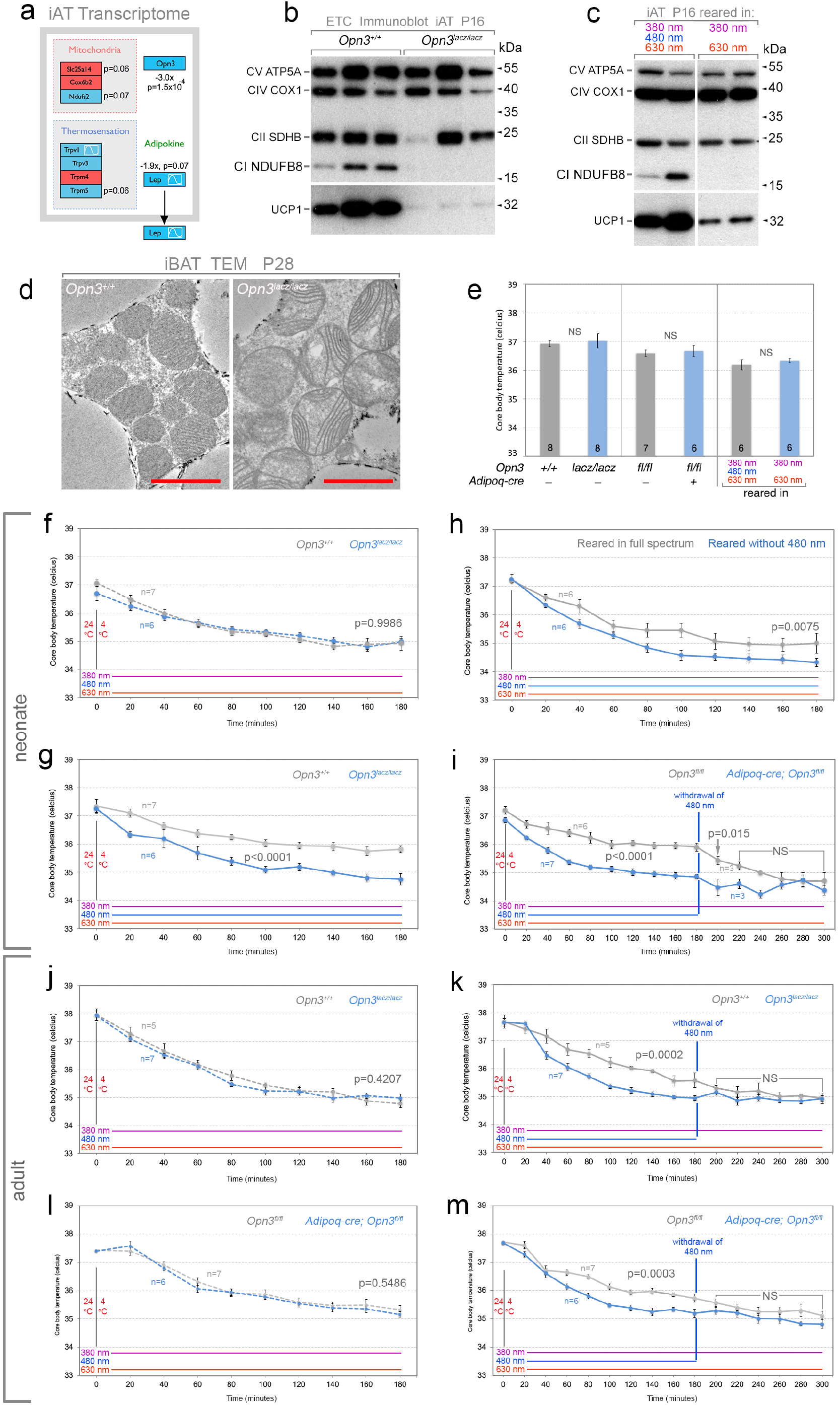
Adipose *Opn3* is required for a normal response to cold stress. (a) Schematic of clustered *Opn3*-dependent transcript changes in iAT for the electron transport chain, as well as the Trp family receptors and Leptin. Red and blue colour coding indicates up- and down-regulated transcripts, respectively. The sine-wave symbol indicates that, according to data within CircaDB, expression of this transcript is diurnally rhythmic in this tissue type. (b, c) Immunoblots detecting multiple components of the electron transport chain (ATP5A, COX1, SDHB, NDUFB8, UCP1) in P16 iAT for *Opn3^+/+^* and *Opn3^lacz/lacz^* (b) and “minus blue” reared mice. NDUFB8 and UCP1 are consistently at lower levels in Opn3 null and “minus blue” reared mice. (d) TEM showing abnormal mitochondrial morphology in the *Opn3* null iBAT at P28. Red scale bar is 2 µm. (e) Core body temperature under ambient conditions of P21 mice that were *Opn3^+/+^* and *Opn3^lacz/lacz^* (left chart), *Opn3^fl/fl^* and *Adipoq-cre; Opn3^fl/fl^* (middle chart) or reared in “minus blue” lighting (right chart). Sample size (n) indicated at base of histogram bar. Significance calculated using Student’s T-test. These measurements were taken 20 minutes before the initiation of the cold stress experiments shown in (f-i). (f-i) Core body temperature assessments over a time course after a 4°C cold challenge for P21-P24 neonatal mice of the indicated genotypes. The lighting conditions used during the cold stress experiments are indicated by the coloured lines above the chart horizontal axis. (f, g) Core body temperature during cold stress in *Opn3^laczlacz^* and control *Opn3^+/+^* in either full spectrum (g, 380 nm + 480 nm + 630 nm) or “minus blue” (f, 380 nm + 630 nm) lighting. These charts show the same cohort of mice. Both *Opn3^laczlacz^* mice and *Opn3^+/+^* mice in “minus blue” show a reduced ability to defend their body temperature. (h) Core body temperature of C57BL/6J mice that were raised from birth either in full spectrum lighting or in the “minus blue” condition. The cold stress experiment was carried out in “full spectrum” light. Mice raised without 480 nm light phenocopy the reduced ability of the *Opn3* null mice to defend their body temperature. (i) As for (g) except the genotypes are *Adipoq-cre; Opn3^fl/fl^* and control *Opn3^fl/fl^*. Also, at minute 180, for one cohort of n=3 *Opn3^fl/fl^* and n=3 *Adipoq-cre; Opn3^fl/fl^* mice, 480 nm light was withdrawn. This resulted in an acute reduction of core body temperature in control mice to that observed in the conditional null. (j, l) As for (f) except adult mice that were *Opn3^laczlacz^* and *Opn3^+/+^* (j) or *Adipoq-cre; Opn3^fl/fl^* and *Opn3^fl/fl^* (l). (k, m) As in (i) except adult *Opn3^laczlacz^* and *Opn3^+/+^* (k) or *Adipoq-cre; Opn3^fl/fl^* and *Opn3^fl/fl^* (m) mice. Sample size (n) labelled on chart. For (f-m), p-values according to ANOVA.

Many features of the *Opn3* germ-line and adipocyte conditional null mice suggested there might be a defect in thermogenesis. UCP1, found at a lower than normal level in both inWAT and iAT of neonatal *Opn3* null mice, has a central role in the thermogenesis response of mice. Furthermore, *Opn3* null mice show low “brite” adipocyte content in inWAT and abnormal mitochondrial morphology in iAT. In neonatal mice, *Leptin*, found at low levels in *Opn3* null mice, is required for establishing the thermogenesis circuit^28^. The TRP channels that cluster in transcriptome analysis of the *Opn3* null, function as thermosensors^29^. There are also strong links between the circadian clock and thermogenesis. When *Bmal1* is conditionally deleted from adipocytes, transcripts for *Ucp1* and *Pparg* upregulate, mitochondrial density is increased and core body temperature is elevated^30^. By contrast, *Per2* null mice under cold stress show reduced transcript levels for *Ucp1* and *Fabp3* (both regulated in the *Opn3* transcriptome analysis, Fig. 5a, b) and are cold sensitive^31^. Finally, NR1D1 (Rev-Erbα) directly regulates *Ucp1* expression in iBAT and *Nr1d1* mutant mice show improved cold tolerance^32^. These data suggest the hypothesis that *Opn3*, through its regulation of the adipocyte clock, may also regulate thermogenesis.

To test this, we performed cold-stress assays on both neonatal and adult mice that were *Opn3* germ line null, adipocyte conditional null, or were raised in “minus blue” conditions. The core body temperature of P21 control and experimental mice within these cohorts was unchanged prior to the cold stress (Fig. 6e). Cold stress assays were performed in lighting conditions that were either “full spectrum” (380 nm, 480 nm and 630 nm) or “minus blue” (380 nm and 630 nm) as a means of assessing whether any changes in thermoregulation were light-dependent. When *Opn3^+/+^* and *Opn3^lacz/lacz^* mice were exposed to 4°C over the course of three hours under “minus blue” lighting conditions, the core body temperature response curves were indistinguishable (Fig. 6f). However, when the same cohorts were assessed a second time under full spectrum lighting, control, *Opn3^+/+^* mice were able to defend their body temperature much more effectively than *Opn3^lacz/lacz^* mice (Fig. 6g). Importantly, because the “minus blue” control and the *Opn3* null mice show a thermogenesis response that is indistinguishable (Fig. 6f), OPN3 function can fully explain the effect of blue light on thermogenesis.

To determine whether neonatal development of the thermogenesis response could be influenced by light, we raised cohorts of C57BL/6J mice from birth in a normal LD cycle of full spectrum or “minus blue” lighting. These were the same conditions used to assess wavelength-dependent neonatal growth (Fig. 1d) and indeed, the “minus blue” reared mice were larger than full spectrum controls at P21 (control, 6.69±0.13 gm, “minus blue” reared, 8.35±0.31 gm, p=0.0007). Despite their elevated body weight, “minus blue” reared mice were less able to defend their body temperature when exposed to a cold stress in full spectrum light (Fig. 6h). This establishes that postnatal development of the thermogenesis response in mice is, surprisingly, dependent on the wavelength of photons to which they are exposed. The correspondence of the thermogenesis defect with that observed in the *Opn3* null, coupled with data indicating that 480 nm light activates OPN3^3^, suggests that development of a normal thermoregulatory circuit is dependent on a light-OPN3 pathway.

To assess whether the defective thermosensory response of the *Opn3* null mice could be attributed to adipocyte OPN3, we repeated the cold stress analysis using cohorts of control *Opn3^fl/fl^* and *Adipoq-cre; Opn3^fl/fl^* mice. In full spectrum lighting, *Adipoq-cre; Opn3^fl/fl^* mice showed a more limited ability than *Opn3^fl/fl^* control mice to defend their body temperature (Fig. 6i). When mice were switched to “minus blue” lighting at 180 minutes of the cold stress experiment, the body temperature of *Opn3^fl/fl^* rapidly changed to the lower body temperature of the conditional null (Fig. 6i). This analysis shows that thermoregulation in neonatal mice is dependent on an adipocyte light-OPN3 pathway that is active both developmentally and acutely. Repetition of cold-stress experiments in adult *Opn3* germ line and adipocyte conditional null mice showed a very similar thermosensory deficit (Fig. 6j-m) in which the absence of blue light, either throughout the cold stress (Fig. 6j,l), or acutely at minute 180 (Fig. 6k,m) could mimic the effect of *Opn3* mutation. These data indicate that adult mice also use an adipocyte light-OPN3 pathway to regulate the use of energy to maintain body temperature.

The analysis presented has assessed the function of Opsin 3^1,3^ (encephalopsin) in the mouse. We show that adipocytes use OPN3 as a light detector in a pathway that regulates photoentrainment of the adipose circadian clock, the composition of adipocyte ECM, the thermosensory response, and growth of neonatal mice. Extraocular photoreception has been described in vertebrates including fish^33,34^ and birds^35^ but there are few examples in mammals^36,37^. The current analysis adds OPN3 to the short list of atypical opsins that act in photoentrainment of circadian clocks. Melanopsin (OPN4) activity can entrain the central circadian clock in the suprachiasmatic nucleus (SCN)^38,10,9^. A second circadian clock system was discovered recently when it was shown that neuropsin (OPN5) is necessary and sufficient for photoentrainment in a retinal circadian clock that functions independently of the OPN4-SCN clock^11,12^. We hypothesize that the OPN3-adipocyte clock represents a third example of a non-visual, opsin-mediated entrainment system, and the first shown to function outside the eye.

A role for the light-OPN3 pathway in regulation of the thermosensory response is likely linked to its role in circadian clock photoentrainment. This is suggested by the many examples of clock gene mutant mice that have thermoregulation deficits and, like the *Opn3* null, show changes in the complement of mitochondrial proteins, such as UCP1, that are required for heat generation^30,31,32^. In some cases it has been shown that genes involved in the thermoregulatory response are direct targets of circadian clock transcription factors^32^. The current analysis indicates that the light-OPN3 pathway is required for development of the thermosensory response^28^. We have shown the light-OPN3 pathway regulates *Leptin*, an adipokine implicated in this aspect of development. Thus, Leptin regulation may explain, at least in part, the action of the light-OPN3 pathway in establishing core body temperature regulation. Since some clock genes can be acutely light-induced, it is also possible that acute modulation of the thermosensory response occurs via acutely regulated clock machinery. We hypothesize that the adipocyte light-OPN3 pathway provides a dynamically responsive, circadian clock-integrated mechanism for matching energy availability to the light-dark cycle.

The adipocyte-specific, OPN3-dependent regulation of heat production also provides an explanation for the overgrowth phenotype observed in the *Opn3* germ line and conditional mutant mice. The generation of heat in endotherms is an adaption for the efficient function of the biochemical processes that constitute normal physiology. In neonatal *Opn3* mutants, we suggest that energy not expended to elevate body temperature can be redirected to the anabolic pathways that underlie growth. Though it might seem that reduced body temperature, and reduced efficiency of anabolism would result in reduced growth, it is likely that under normal conditions, *Opn3* neonatal mice maintain their body temperature by huddling with the dam and their littermates. It has been documented that individual new-born rodents with reduced heat production capacity can acquire heat energy from the dam and from littermates^39^. The absence of a significant core body temperature change in *Opn3* mutant mice under ambient conditions further suggests that only when *Opn3* mutant mice are isolated and cold-stressed is the heat generation abnormality observable.

In sum, we have found that blue light significantly regulates neonatal growth and metabolism in mice. Unexpectedly, we find this effect is mediated by direct photoreception in adipose tissue, via an ‘orphan’ opsin, OPN3. Direct photosensitivity of isolated adipose tissue is demonstrated by its ability to synchronize its circadian clock to light-dark cycles, a phenomenon which requires OPN3 function. Loss of OPN3 results in alteration of numerous metabolic pathways, including modified cellular composition of fat, reduced NAD levels, and defective thermogenesis under cold challenge. While all these effects are light-dependent, it remains possible that OPN3 also has light-independent basal functions. Further, it is not yet clear which OPN3 effects are primary; does disruption of the circadian clock cause the observed metabolic changes, or does altered adipocyte metabolism negatively impact the clock? Or, are both involved in a more complicated feedback loop? Further study of clock mutant mice under similar physiologic conditions may help resolve these questions.

OPN3 is highly conserved across species and is expressed in human adipocytes. If the light-OPN3 adipocyte pathway exists in humans, there are potentially broad implications for human health. Our modern lifestyle subjects us to unnatural lighting spectra, exposure to light at night, shift work and jet-lag, all of which result in metabolic disruption^40,41^ ^42,43^. Recent studies have suggested that light exposure directly affects metabolic function in humans^44^. Based on the current findings, it is quite likely that insufficient or arrhythmic stimulation of the light-OPN3 adipocyte pathway is part of an explanation for the prevalence of metabolic deregulation in the industrialized nations where unnatural lighting patterns have become the norm.

## Figures and Legends

**Figure S1.**
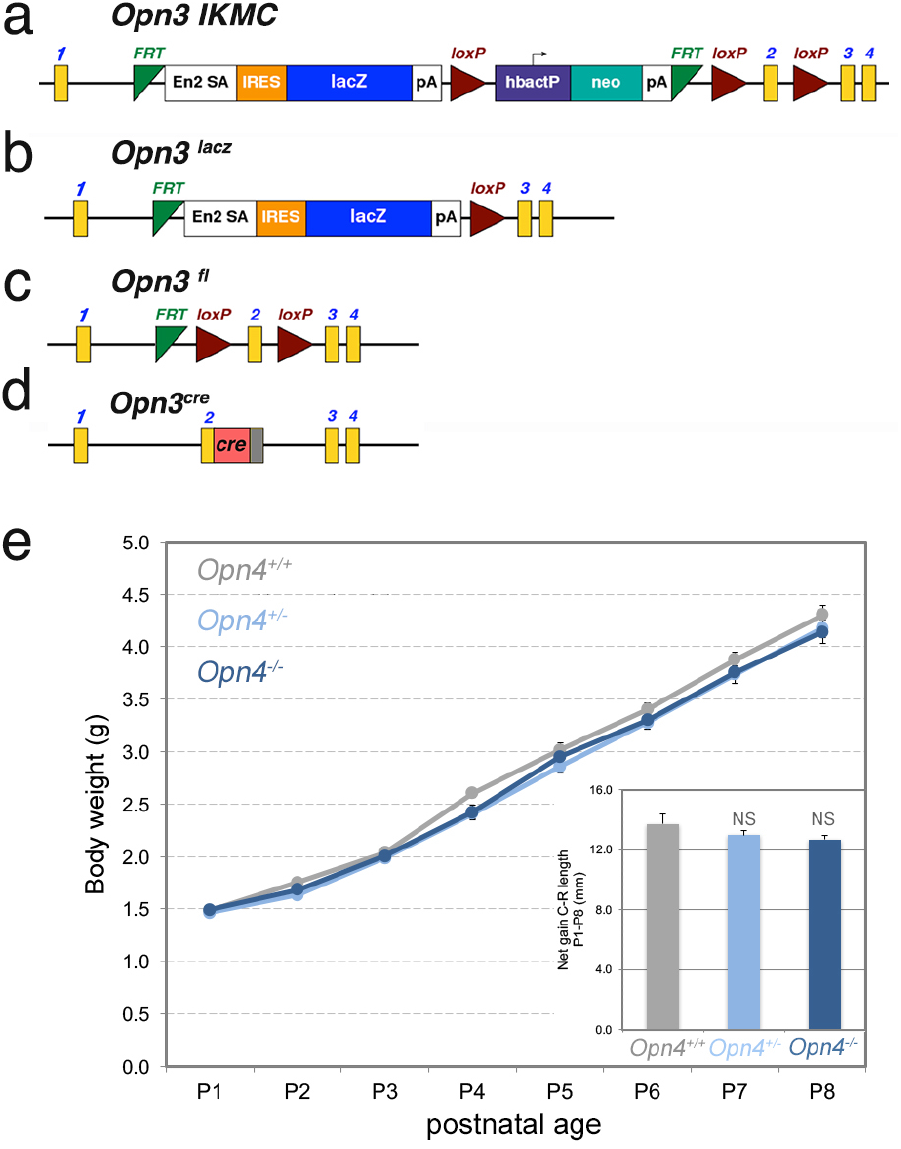
*Opn3* alleles and *Opn4* null neonatal growth. (a-d) Schematics of the *Opn3* alleles used in this study. (a) The *Opn3* allele as targeted in ES cells by the International Knockout Mouse Consortium. Exons are numbered yellow boxes. *FRT*, FLP recombinase site-specific recombination sites. En2 SA, *Engrailed 2* splice acceptor. IRES, internal ribosome entry sequence. *Lacz*, β-galactosidase open reading frame. pA, polyadenylation signal. loxP, cre recombinase site specific recombination sequences. hbactP, human β-actin promoter. Right-facing arrow indicates the start point of transcription for hbactP. neo, the neomycin resistance gene. (b) The *Opn3^lacz^* allele generated after germ-line recombination at the *loxP* sites. (c) the *Opn3^fl^* allele generated after germ-line recombination at the *FRT* sites. The *Opn3^cre^*allele was generated by inserting the cre recombinase open reading frame into exon 2 using CRISPR/Cas9. (e) Body weight and crown-to rump length (inset) for *Opn4^+/+^* (grey, n=6), *Opn4^+/^-* (light blue, n=23) and *Opn4^-/-^*(dark blue, n=10) mice over the P1 to P8 time-course. According to ANOVA, no statistical differences between genotypes were identified.

**Figure S2.**
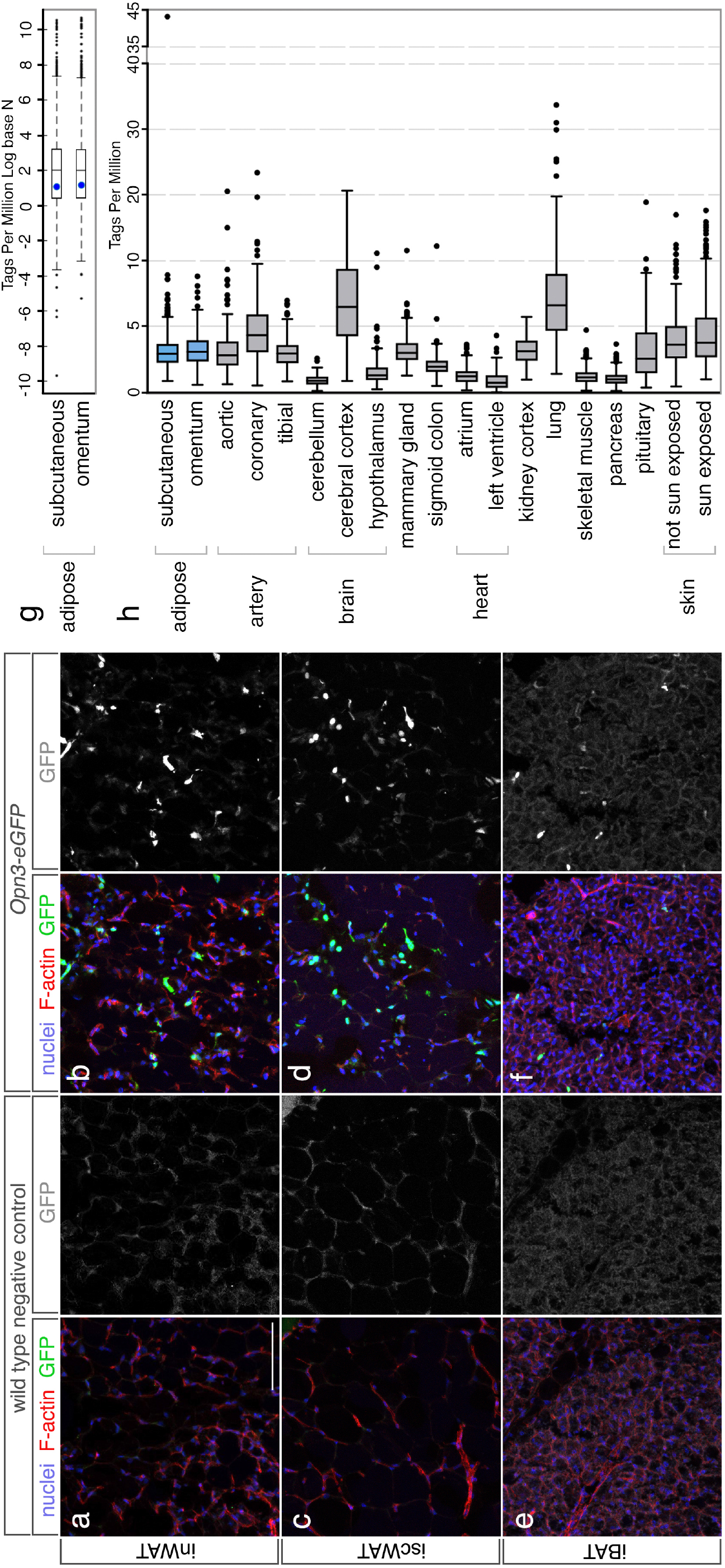
Expression of *Opn3-eGFP* reporter in adipose tissue. (a-f) Panels show cryosections of adipose tissue from P16 mice labeled with Hoechst33258 to detect nuclei (blue), with fluorochrome-conjugated Phalloidin to detect F-actin (red), and with anti-GFP antibodies (green, gray). (a, b) Inguinal white adipose tissue (inWAT), (c, d) interscapular white adipose tissue (iscWAT), (e, f) interscapular brown adipose tissue (iBAT). For clarity, the green channel detecting GFP is shown in grayscale. White, 100 µm scale bar in (a) applies to all immunofluorescence panels. (g) Box and whiskers plot showing Tags Per MIllion (TPM)(Log base N) sequencing reads for the *OPN3* transcript (blue dot) versus all other transcripts in human subcutaneous and omental adipose tissue. The box defines the interquartile data range (IQR, the difference between the 25^th^ and 75^th^ percentiles), and the line within the box, the median. Box plot whiskers are plotted according to the Tukey method. The *OPN3* transcript is detected at 2.9 TPM for subcutaneous adipose and 3.1 TPM for omental adipose. The median value for all transcripts in these adipose tissues is 7.6 TPM for subcutaneous and 7.0 for omental. (h) Box and whiskers plot showing Tags Per MIllion sequencing reads for human *OPN3* transcript in the indicated tissues. Subcutaneous and omental adipose tissue median *OPN3* expression is in the mid-range of expression values.

**Figure S3.**
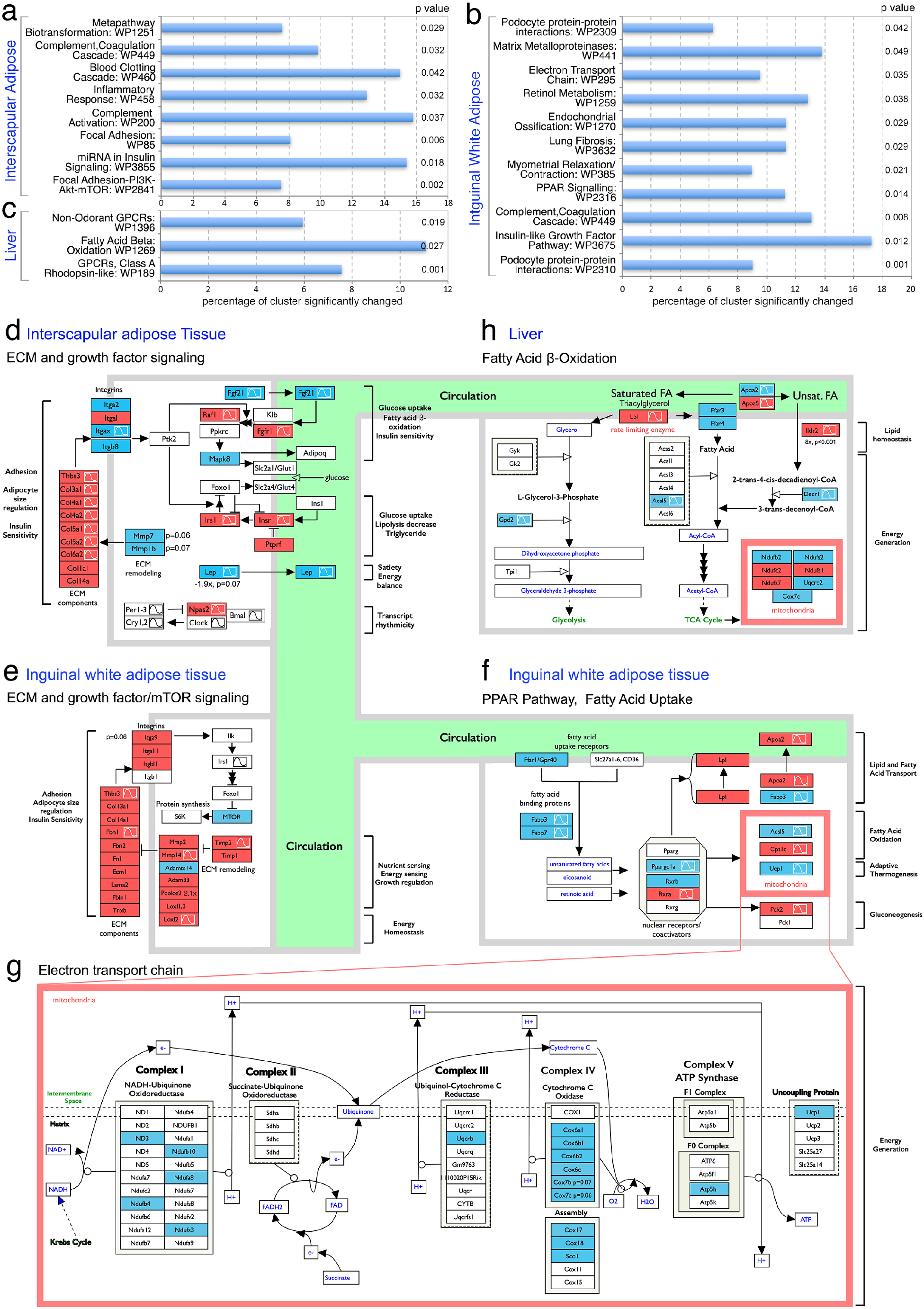
Transcriptome analysis of *Opn3* null mice. (a-c) Charts show, for interscapular adipose tissue (a), inguinal white adipose tissue (b) and liver (c), the WikiPathways (text labels and WP numbers) in which there was significant clustering (according to Z-score) of significantly changed transcripts in *Opn3^+/+^* versus *Opn3^lacz/lacz^* tissue. The chart horizontal axis shows the percentage of significantly changed transcripts for a given pathway. Fisher’s exact test p-values are listed on the right of the chart. (d-h) Panels show schematically, in the context of known pathways, *Opn3*-dependent changes in transcripts for the indicated genes in iAT (d) inWAT (e-g) and liver (h). Each box represents a transcript that is significantly (p<0.05) up-regulated (red) or down-regulated (blue) in the *Opn3* null. For four transcripts (*Mmp7, Mmp1b, Lep* (d) and *Itga9* (e)) where the p-value did not quite reach p<0.05, the p-value is listed. The sine-wave symbol indicates that, according to data within CircaDB, expression of this transcript is diurnally rhythmic in this tissue type. The circulation is modelled by the green area. All panels are modified versions of Wikipathway schematics for Focal Adhesion-PI3K-mTOR Signaling pathway (WP2841, (d, e)), for the PPAR Signaling Pathway (WP2316, (f)), for the Electron Transport Chain (WP295, (g)) and for the Fatty Acid β-Oxidation Pathway (WP1269, (h)).

**Figure S4.**
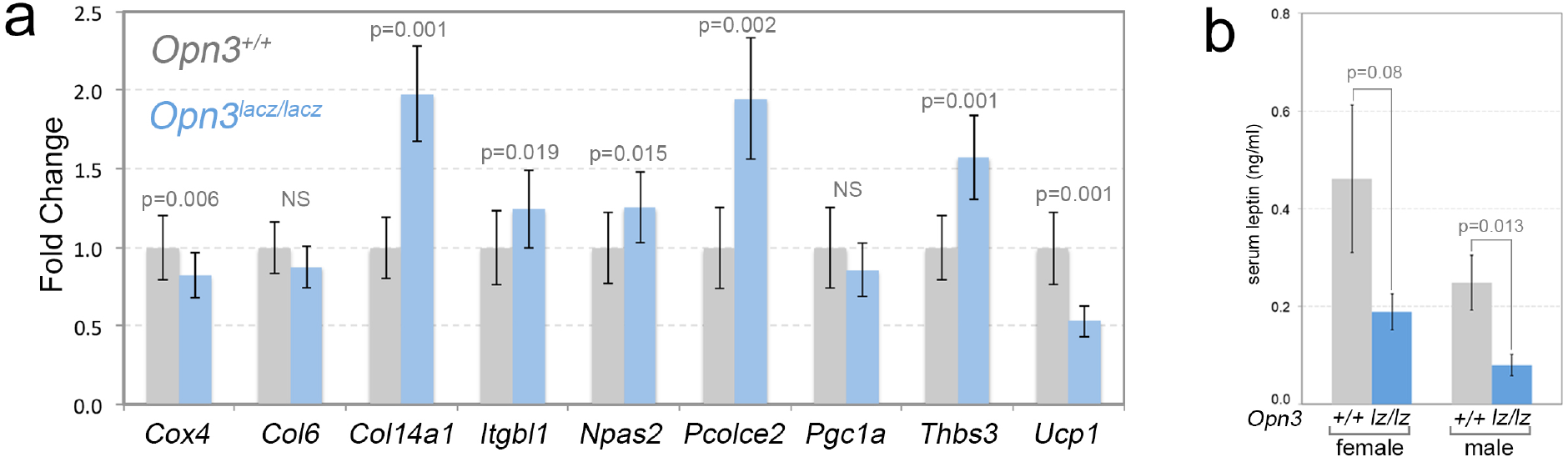
Transcriptome validation and Leptin in *Opn3* null mice. (a) qPCR assessment of relative expression levels for the indicated transcripts in inWAT of P16 *Opn3^+/+^* and *Opn3^lacz/lacz^* mice. n=3 for each genotype. (b) Chart showing serum leptin levels in adult female (control, n=8, *Opn3^lacz/lacz^*, n=8) and male (control, n=7, *Opn3^lacz/lacz^*, n=8) mice. p-values calculated using Student’s T-test.

## Materials and Methods

### Mice

Animals were housed in a pathogen-free vivarium in accordance with institutional policies. Genetically modified mice used in this study were: *B6;FVB-Tg(Adipoq-cre)1Evdr/J*^1^ (Jax stock #010803), *Ai14*^2^ (Jax stock #007914), *Opn4*^3^, *Tg(Opn3-EGFP)JY3Gsat* (MMRRC stock number 030727-UCD) and *Opn3^tm2a(EUCOMM)Wtsi^* mice that were generated from C57BL/6N ES cells obtained from EUCOMM (ES clone ID: EPD0197_3_E01). The ES cells harbour a genetic modification wherein the lacz-Neomycin cassette is flanked by FRT sites and a *loxp* site separates lacz from the neomycin coding region. *Loxp* sites also flank exon2 of *Opn3* allowing multiple mouse lines that can serve as reporter nulls, conditional floxed and null mice. The *Opn3^Lacz^* reporter-null line was created by crossing *Opn3^tm2a(EUCOMM)Wtsi^* mice to *FVB/N-Tg(EIIa-cre)C5379Lmgd/J mice*^4^ (Jax stock #003314). *Opn3^fl/fl^* line was created by crossing the *Opn3^tm2a(EUCOMM)Wtsi^* mice to *129S4/SvJaeSor-Gt(ROSA)26Sor^tm1(FLP1)Dym^/J*^5^ (Jax stock #003946) to remove the *lacZ* cassette. This means that *Opn3^lacz^* mice are of mixed C57Bl6/6N, FVB/N background and that *Adipoq-cre; Opn3^fl^* mice are of mixed C57Bl6/6N, 129S4/Sv, B6;FVB background. Littermate control animals were used for all experiments with the exception of C57BL/6J mice reared under different lighting conditions.

The *Opn3^cre^* was generated in-house using CRISPR-Cas9 technology. Four gRNAs that target exon 2 of *Opn3* were selected to knock in the Cre cassette. Plasmids containing the gRNA sequence were transfected into MK4 cells (an in-house mouse cell line representing induced metanephric mesenchyme undergoing epithelial conversion). The editing efficiency of gRNA was determined by T7E1 assay of PCR products of the target region amplified from genomic DNA of transfected MK4 cells. The sequence of the gRNA that was subsequently used for the transfection is <u>TACCGTGGACTGGAGATCCA.</u> Sanger sequencing was performed to validate the knock-in sequence of founder mice.

Mice were placed on normal chow diet (NCD: 29% Protein, 13% Fat and 58% Carbohydrate kcal; LAB Diet #5010) ad libitum with free access to water. Serum for leptin was collected by tail bleed after a 6 hour daytime food withdrawal at age 16 weeks.

### Genotyping

Primer sequences for genotyping the *Opn3^fl^* and *Opn3^lacz^* alleles are:

F1: ACCCAGGCTTCTTTTGGTCT; R1: AGAGTCGTTGGCATCCTTGG; F2: ACTATCCCGACCGCCTTACT; R2: GAACTGATGGCGAGCTCAGA

F1-R1 gives a 1191 bp wild-type band; F1-R1 gives a 1231 bp band from the *Opn3^fl^* allele. F2-R2 gives a 701 bp band from the *Opn3^lacz^* reporter null allele. Primer sequences for genotyping the *Opn3^cre^* allele are: Opn3creF1:TGCTGGCCTATGAACGTTATATCC;

Opn3creR2: CACTCGTTGCATCGACCGGTAATGC. These give a 390 bp band.

### Lighting conditions

Animals were housed in standard fluorescent lighting (photon flux 1.62×10^15^ photons/cm^2^/sec) on a 12L:12D cycle except where noted. For full spectrum lighting, LEDs were used to yield a comparable total photon flux of 1.68×10^15^ photons/cm^2^/sec. Spectral and photon flux information for LED lighting: near-violet (λ_max_=380 nm, 4.23×10^14^ photons/cm^2^/sec in 370-400 nm range), blue (λ_max_=480 nm, 5.36×10^15^ photons/cm^2^/sec in 430-530 nm range), green (λ_max_=530 nm, 5.82×10^15^ photons/cm^2^/sec in 480-600 nm range) and red (λ_max_=630 nm, 1.93×10^15^ photons/cm^2^/sec in 590-660 nm range). For wavelength restricted growth assessment, C57BL/6J animals were housed in 12L:12D cycle starting late gestation (embryonic day E16) either with blue (480 nm and 380nm LEDs) or without blue (380 nm LEDs) lighting.

**Measurement of bioluminescence from tissues of *Per2^Luciferase^* mice**.*Per2::Luciferase* mice^6^ were euthanized by CO_2_ asphyxiation, and scapular and inguinal fat pads were quickly dissected and placed into cold Hank’s Balanced Salt Solution (HBSS, Gibco). WAT was dissected as ∼3 mm pieces and cultured on cell culture inserts (Millipore PICMORG50) in 35 mm diameter dishes in DMEM containing B27 serum-free supplement (Life Technologies), 352.5 µg/ml sodium bicarbonate (Life Technologies), 10 mM HEPES (Life Technologies), 25 units/ml penicillin, 25 µg/ml streptomycin (Life Technologies), 10 µM 9-*cis*-retinaldehyde (Sigma-Aldrich), and 100 µM luciferin potassium salt (Biosynth). Dishes were sealed with sterile vacuum grease and placed in a Lumicycle photomultiplier tube (PMT) system (Actimetrics). All tissue was maintained at 36°C. Bioluminescence data was measured and exported as raw counts (counts/second) by Lumicycle Analysis software (Actimetrics). For determination of peak/trough amplitude, raw counts were normalized to the lowest count for each trace from individual tissues. The trough value was compared to the peak value of luminescence between 1.5 and 2.5 days after culture.

For photoentrainment of tissues, tissues were dissected as described above but then placed into 24-hour clock motorized photoentrainment devices for 5 days^7^. Pairs of tissues experienced 5 days of anti-phasic 9h:15h light:dark cycles, and tissues were then transferred, in darkness, to a Lumicycle PMT system. The light for light:dark cycles was produced by 417-nm, 475-nm, and 530-nm LEDs for a combined output of 4W/m^2^. Bioluminescence traces were analysed for period and phase measurements by the sine wave best-fitted to at least 3 days of oscillations using Lumicycle Analysis software.

### Immunohistochemistry and tissue processing

Animals were anesthetized under isoflurane and sacrificed by cervical dislocation. Adipose tissue depots (interscapular adipose tissue complex and inguinal WAT) were harvested and fixed in ice cold 10 % zinc formalin for 1 hour at 4°C. After washing in PBS, adipose tissue samples were prepared for cryosectioning as described previously^8^. Gelatin-embedded tissues were sectioned at 16 μm in a cryostat and labelled with primary antibodies as previously described^8^. Rabbit antibodies to GFP (ab13970, 1 in 500), Collagen VI (COL6, ab6588, 1 in 500), COX4 (Genetex, GTX114330, 1 in 400, cold acetone:methanol, 1:1, post-fixed, 10 min), and UCP1 (ab10983, 1 in 500), were purchased from Abcam. Alexa 647 conjugated isolectin (1 in 300) and Alexa 546 conjugated F-actin were purchased from Invitrogen. Alexa 488 conjugated secondary antibodies (1 in 300) were purchased from Jackson ImmunoResearch.

### X-Gal staining

For X-Gal labelling, tissue samples were fixed in X-Gal fixative (1% formaldehyde, 0.2% glutaraldehyde, 2 mM MgCl_2_, 5 mM EGTA, and 0.01% Nonidet P-40) for two hours at room temperature. Tissues were cryosectioned as described above and then labelled with X-Gal. The reaction was monitored closely and stopped when background started to appear in control (wild-type) tissues. Following two washes in PBS, cryosections were imaged using a bright field microscope.

### Hemotoxylin labelling and cell-size quantification

Gelatin-embedded frozen sections of inguinal WAT (as described above) were stained with hematoxylin and imaged under bright-field. Samples were imaged with a rhodamine filter to assess adipocyte size distribution. Using the free hand selection tool on ImageJ, adipocytes were outlined and the area measured in μm^2^. Cell size distribution was determined by quantifying 60 cells from at least 10 regions, for a total of approximately 600 cells per animal. Cell areas were binned into 200 μm^2^ intervals and the frequency of total cells (%) charted for each interval.

### Western Blotting

Western blots were performed using standard protocols. Adipose tissue lysates were made in NP40 lysis buffer: 150 mM NaCl, 1% NP40, 50 mM Tris 8.0 with phosphatase inhibitors. Lysates were prepared by sonication and the lysates were separated from overlaying fat layer by three rounds of centrifugation. After BCA method of protein quantification, lysates were boiled in Laemmli sample buffer (4% SDS, 20% glycerol, 10% 2-mercaptoethanol, 0.004% bromophenol blue and 0.125 M Tris HCl, pH 6.8). Blots were incubated with OxPhos antibodies (Thermofisher 45-8099 1:1000) and UCP-1 (Abcam ab10983). HRP-conjugated secondary antibodies were used at 1:5000 dilution and detected by enhanced chemiluminescence (ThermoFisher Scientific).

### Microarray analysis

Interscapular adipose tissue complex and inguinal white adipose tissue from P16 mice were harvested at one hour after lights on (ZT1) and snap frozen on dry ice. Tissue pieces were homogenized in TRIzol (TriReagent Invitrogen) using RNAse-free Zirconium oxide beads (2.0 mm) in a TissueLyser II (Qiagen). Phase separation was achieved using chloroform and RNA in the aqueous phase was precipitated using ethanol. RNA was purified by column method using GeneJET RNA purification kit (ThermoFisher Scientific #K0732) and eluted into RNAse-free water. RNA quality was assessed using the Agilent 2100 Bioanalyzer and an RNA-integrity number cut-off of 7 was applied for selecting samples for microarray assay. RNA from biological triplicates were submitted for microarray assay (ClariomD, Affymetrix) to the Technology Center for Genomics and Bioinformatics, University of California, Los Angeles.

Data analysis including normalization, gene expression changes and gene-enrichment analysis was performed using AltAnalyze, developed by Nathan Salomonis at Cincinnati Children’s Hospital Medical Center. AltAnalyze uses the robust multi-array average method of normalization. Briefly, the raw intensity values are background corrected, log2 transformed and then quantile normalized. Next, a linear model is fit to the normalized data to obtain an expression measure for each probe set on each array. Gene expression changes greater than 1.1 fold were calculated using unpaired t-test, where a p-value <0.05 was used as a cut-off.

### Quantitative RT-PCR

Inguinal adipose depot was harvested at indicated times of day (ZT1, one hour after lights ON and ZT17, 5 hours after lights OFF) from P16 mouse pups. Samples at ZT17 were harvested under dim red light. Snap frozen tissue was homogenized and processed for RNA as described above. RNA was treated with RNase-free DNase I (ThermoFisher Scientific #EN0521) and cDNA was synthesized using Verso cDNA synthesis kit (ThermoFisher Scientific AB1453/B). Quantitative RT-PCR was performed with Radiant™ SYBR Green Lo-ROX qPCR mix (Alkali Scientific Inc.) in a ThermoFisher QuantStudio 6 Flex Real-Time PCR system. Primer information for quantitative PCR is included in the Table. Relative expression software tool (REST^©^)^9^ was used for statistical analysis (fixed pair-wise randomization reallocation test) of qPCR results using three normalizing genes (*Hprt, Rplp0, Hmbs*).

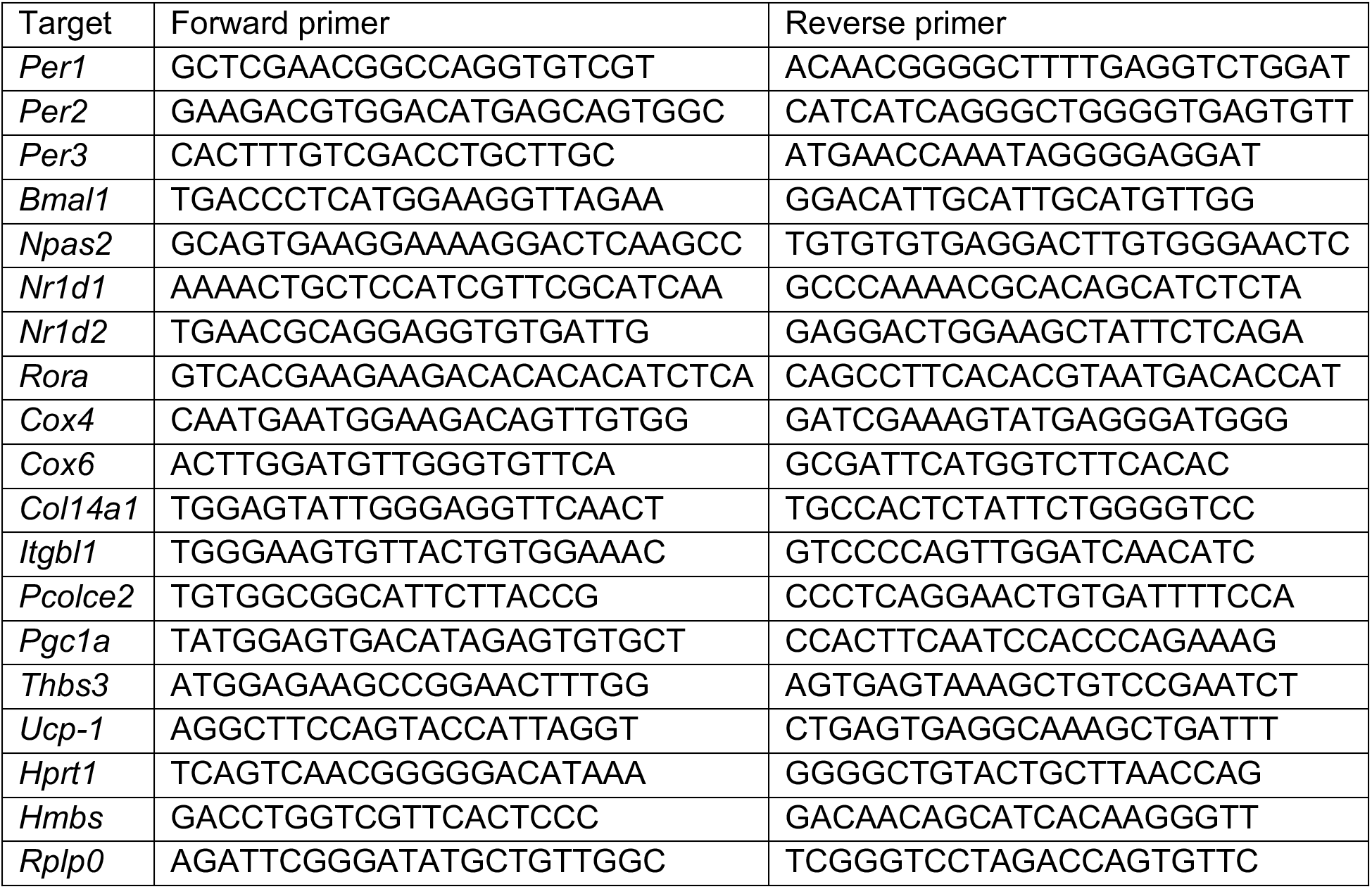

### Transmission Electron Microscopy

Freshly dissected adipose tissues were collected and 1 mm samples from approximately similar areas were fixed in 2% glutaraldehyde, 1% paraformaldehyde in PBS for 1 hour at room temperature before processing and sectioning for transmission electron microscopy as described before^10^.

### NAD/NADH Quantification

NAD levels were measured using NAD/NADH assay kit from Abcam (ab65348). Briefly, tissues samples (inguinal adipose tissue and liver) from P16 mouse pups were snap frozen in liquid nitrogen, homogenized in NADH/NAD extraction buffer and filtered through a 10kD spin column (ab93349) to remove enzymes. Assay procedure was followed per kit instructions and levels of NADH and NAD+ were determined normalized to tissue weight.

### Thermoregulation assay

Acute cold temperature challenge was performed on control and experimental mice from *Opn3* reporter null (*Opn3^+/+^* and *Opn3^lacz/lacz^*) and adipose tissue conditional deletion of *Opn3* (*Opn3^fl/fl^* and *Adipoqcre; Opn3^fl/fl^*) mouse lines. In addition, C57BL/6J mice reared under wavelength restriction (with or without blue, as described previously) were subjected to this assay. Littermates housed with mother were separated from their home cage and individually housed in a home-built lighting chamber situated in an electronically monitored 4°C cold room. While the mouse was conscious, body temperature was measured rectally with a RET-3 Microprobe Thermometer (Kent Scientific) every 20 minutes for the duration of the assay. Food and water were available *ad libitum*. The thermo probe operator was blinded to mouse genotype and prior temperature measurements throughout the study. At the end of the 3-hour cold stress, the mice were returned to their home cage. For the 3-hour cold challenge at postnatal day 21, mice were subjected to near-violet (380 nm) and red (630 nm) wavelengths in the light chamber. A second 3-hour cold challenge was conducted at postnatal day 24 where the mice were subjected to near-violet (380 nm), blue (480 nm) and red (630 nm) wavelengths. The exception being Fig 6g, where the study was extended by 2 hours following withdrawal of 480 nm wavelength. The order of cage placement was randomized at this time, such that the thermo probe operator remained blinded.

### Data analysis

Statistical analyses were performed using GraphPad Prism version 4.00 (GraphPad Software), Microsoft Excel and Sigma plot (Figure 3). Growth charts were analysed by two-way ANOVA followed by *t*-test for comparison at each time-point. Two-tailed distribution, two-sample unequal variance *t*-test was used to determine the statistical significance between two independent groups.

Author Contribution
SV, GN, EB: Experimental design and analysis, manuscript preparation. ANS, JZ, JAM, KXZ, BAU, YO, VB, EB, KM, M-TN, SAG, GW, RS, XM, SR: Experimental execution and analysis. NTP, MB: Electronic device design and construction. JBH: Supervision of bioinformatics analysis. TN, RNVG, RAL: Project leadership, experimental design and manuscript preparation.

## References

1. Blackshaw, S. & Snyder, S. H. Encephalopsin: a novel mammalian extraretinal opsin discretely localized in the brain. J Neurosci 19, 3681–3690 (1999).

2. Arendt, D. Ciliary Photoreceptors with a Vertebrate-Type Opsin in an Invertebrate Brain. Science (80-.). 306, 869–871 (2004).

3. Koyanagi, M., Takada, E., Nagata, T., Tsukamoto, H. & Terakita, A. Homologs of vertebrate Opn3 potentially serve as a light sensor in nonphotoreceptive tissue. Proc Natl Acad Sci U S A 110, 4998–5003 (2013).

4. Sugihara, T., Nagata, T., Mason, B., Koyanagi, M. & Terakita, A. Absorption characteristics of vertebrate non-visual opsin, Opn3. PLoS One 11, (2016).

5. Regard, J. B., Sato, I. T. & Coughlin, S. R. Anatomical Profiling of G Protein-Coupled Receptor Expression. Cell 135, 561–571 (2008).

6. Hardy, O. T. et al. Body mass index-independent inflammation in omental adipose tissue associated with insulin resistance in morbid obesity. Surg. Obes. Relat. Dis. 7, 60–7 (2011).

7. Eguchi, J. et al. Transcriptional control of adipose lipid handling by IRF4. Cell Metab 13, 249–259 (2011).

8. Rosenwald, M., Perdikari, A., Rülicke, T. & Wolfrum, C. Bi-directional interconversion of brite and white adipocytes. Nat. Cell Biol. 15, 659–67 (2013).

9. Berson, D. M., Dunn, F. A. & Takao, M. Phototransduction By Retinal Ganglion Cells That Set The Circadian Clock. Science (80-.). 295, 1070–1073 (2002).

10. Hattar, S., Liao, H. W., Takao, M., Berson, D. M. & Yau, K. W. Melanopsin-containing retinal ganglion cells: architecture, projections, and intrinsic photosensitivity. Science (80-.). 295, 1065–1070 (2002).

11. Buhr, E. D. & Van Gelder, R. N. Local photic entrainment of the retinal circadian oscillator in the absence of rods, cones, and melanopsin. Proc Natl Acad Sci U S A 111, 8625–8630 (2014).

12. Buhr, E. D. et al. Neuropsin (OPN5)-mediated photoentrainment of local circadian oscillators in mammalian retina and cornea. Proc Natl Acad Sci U S A 112, 13093–13098 (2015).

13. Yoo, S. H. et al. PERIOD2::LUCIFERASE real-time reporting of circadian dynamics reveals persistent circadian oscillations in mouse peripheral tissues. Proc Natl Acad Sci U S A 101, 5339–5346 (2004).

14. Izumo, M. et al. Differential effects of light and feeding on circadian organization of peripheral clocks in a forebrain Bmal1 mutant. Elife 3, (2014).

15. Salomonis, N. AltAnalyze –­ An optimized platform for RNA-seq splicing and domain-level analyses. in Proceedings –­ 2012 IEEE 2nd Conference on Healthcare Informatics, Imaging and Systems Biology, HISB 2012 113 (2012). doi:10.1109/HISB.2012.38

16. Emig, D. et al. AltAnalyze and DomainGraph: Analyzing and visualizing exon expression data. Nucleic Acids Res. 38, (2010).

17. Khan, T. et al. Metabolic dysregulation and adipose tissue fibrosis: role of collagen VI. Mol. Cell. Biol. 29, 1575–1591 (2009).

18. Spencer, M. et al. Adipose tissue extracellular matrix and vascular abnormalities in obesity and insulin resistance. J. Clin. Endocrinol. Metab. 96, E1990–8 (2011).

19. Divoux, A. & Clément, K. Architecture and the extracellular matrix: The still unappreciated components of the adipose tissue. Obes. Rev. 12, (2011).

20. Mariman, E. C. M. & Wang, P. Adipocyte extracellular matrix composition, dynamics and role in obesity. Cell. Mol. Life Sci. 67, 1277–1292 (2010).

21. Pizarro, A., Hayer, K., Lahens, N. F. & Hogenesch, J. B. CircaDB: a database of mammalian circadian gene expression profiles. Nucleic Acids Res 41, D1009–13 (2013).

22. Bray, M. S. et al. Disruption of the circadian clock within the cardiomyocyte influences myocardial contractile function, metabolism, and gene expression. Am. J. Physiol. Heart Circ. Physiol. 294, 1036–1047 (2008).

23. Liu, C., Li, S., Liu, T., Borjigin, J. & Lin, J. D. Transcriptional coactivator PGC-1α integrates the mammalian clock and energy metabolism. Nature 447, 477–481 (2007).

24. Fan, W. & Evans, R. PPARs and ERRs: Molecular mediators of mitochondrial metabolism. Current Opinion in Cell Biology 33, 49–54 (2015).

25. Wu, J. et al. Beige adipocytes are a distinct type of thermogenic fat cell in mouse and human. Cell 150, 366–376 (2012).

26. Cousin, B. et al. Occurrence of brown adipocytes in rat white adipose tissue: molecular and morphological characterization. J. Cell Sci. 103, 931–942 (1992).

27. Peek, C. B. et al. Circadian clock NAD+ cycle drives mitochondrial oxidative metabolism in mice. Science (80-.). 342, (2013).

28. Zeltser, L. M. Developmental influences on circuits programming susceptibility to obesity. Frontiers in Neuroendocrinology 39, 17–27 (2015).

29. Vriens, J., Nilius, B. & Voets, T. Peripheral thermosensation in mammals. Nature Reviews Neuroscience 15, 573–589 (2014).

30. Nam, D. et al. The adipocyte clock controls brown adipogenesis via TGF-beta/BMP signaling pathway. J Cell Sci (2015). doi:10.1242/jcs.167643

31. Chappuis, S. et al. Role of the circadian clock gene Per2 in adaptation to cold temperature. Mol Metab 2, 184–193 (2013).

32. Gerhart-Hines, Z. et al. The nuclear receptor Rev-erbalpha controls circadian thermogenic plasticity. Nature 503, 410–413 (2013).

33. Kojima, D. & Fukada, Y. Non-visual photoreception by a variety of vertebrate opsins. Novartis Found Symp 224, 265–282 (1999).

34. Sato, K. et al. Two UV-sensitive photoreceptor proteins, Opn5m and Opn5m2 in ray-finned fish with distinct molecular properties and broad distribution in the retina and brain. PLoS One 11, (2016).

35. Nakane, Y. et al. A mammalian neural tissue opsin (Opsin 5) is a deep brain photoreceptor in birds. Proc. Natl. Acad. Sci. 107, 15264–15268 (2010).

36. Sikka, G., Hori, D., Pandey, D., Barreto, S. & Berkowitz, D. Opsin 3 and 4 mediate light-dependent vasorelaxation: Therapeutic targets in pulmonary hypertension. Crit. Care Med. 44, 138 (2016).

37. Sikka, G. et al. Melanopsin mediates light-dependent relaxation in blood vessels. Proc. Natl. Acad. Sci. 111, 17977–17982 (2014).

38. Stetson, M. H. & Watson-Whitmyre, M. Nucleus suprachiasmaticus: the biological clock in the hamster? Science 191, 197–9 (1976).

39. Haig, D. Huddling: Brown Fat, Genomic Imprinting and the Warm Inner Glow. Current Biology 18, (2008).

40. Fonken, L. K. & Nelson, R. J. The effects of light at night on circadian clocks and metabolism. Endocrine Reviews 35, 648–670 (2014).

41. Opperhuizen, A. L. et al. Light at night acutely impairs glucose tolerance in a time-, intensity- and wavelength-dependent manner in rats. Diabetologia 1–11 (2017). doi:10.1007/s00125-017-4262-y

42. Fonken, L. K., Aubrecht, T. G., Meléndez-Fernández, O. H., Weil, Z. M. & Nelson, R. J. Dim Light at Night Disrupts Molecular Circadian Rhythms and Increases Body Weight. J. Biol. Rhythms 28, 262–271 (2013).

43. Laermans, J. & Depoortere, I. Chronobesity: Role of the circadian system in the obesity epidemic. Obes. Rev. 17, 108–125 (2016).

44. Cheung, I. N. et al. Morning and Evening Blue-Enriched Light Exposure Alters Metabolic Function in Normal Weight Adults. PLoS One 11, (2016).

## References

1. Eguchi, J. et al. Transcriptional control of adipose lipid handling by IRF4. Cell Metab. 13, 249–259 (2011).

2. Madisen, L. et al. A robust and high-throughput Cre reporting and characterization system for the whole mouse brain. Nat Neurosci 13, 133–140 (2010).

3. Panda, S. et al. Melanopsin is required for non-image-forming photic responses in blind mice. Science (80-.). 301, 525–527 (2003).

4. Lakso, M. et al. Efficient in vivo manipulation of mouse genomic sequences at the zygote stage. Proc. Natl. Acad. Sci. U. S. A. 93, 5860–5 (1996).

5. Henrich, V. C. et al. Widespread recombinase expression using FLPeR (flipper) mice. Genesis 28, 106–110 (2000).

6. Yoo, S. H. et al. PERIOD2::LUCIFERASE real-time reporting of circadian dynamics reveals persistent circadian oscillations in mouse peripheral tissues. Proc Natl Acad Sci U S A 101, 5339–5346 (2004).

7. Buhr, E. D. & Van Gelder, R. N. Local photic entrainment of the retinal circadian oscillator in the absence of rods, cones, and melanopsin. Proc Natl Acad Sci U S A 111, 8625–8630 (2014).

8. Berry, R. et al. Imaging of adipose tissue. Methods Enzymol. 537, 47–73 (2014).

9. Pfaffl, M. W., Horgan, G. W. & Dempfle, L. Relative expression software tool (REST) for group-wise comparison and statistical analysis of relative expression results in real-time PCR. Nucleic Acids Res. 30, e36 (2002).

10. Cinti S, Zingaretti MC, Cancello R, Ceresi E, F. P. Morphologic techniques for the study of brown adipose tissue and white adipose tissue. Methods Mol. Biol. 155, 21–51 (2001).

